# OTUB1 non-catalytically regulates the stability of the E2 ubiquitin conjugating enzyme UBE2E1

**DOI:** 10.1101/359414

**Authors:** Nagesh Pasupala, Marie E. Morrow, Lauren T. Que, Barbara A. Malynn, Averil Ma, Cynthia Wolberger

**Author notes:** These authors contributed equally (Nagesh Pasupala, Marie E. Morrow).

## Abstract

OTUB1 is a deubiquitinating enzyme that cleaves K48-linked polyubiquitin chains and also regulates ubiquitin signaling through a unique, non-catalytic mechanism. OTUB1 binds to a subset of E2 ubiquitin conjugating enzymes and inhibits their activity by trapping the E2~ubiquitin thioester and preventing ubiquitin transfer. The same set of E2s stimulate the deubiquitinating activity of OTUB1 when the E2 is not charged with ubiquitin. Previous studies have shown that, in cells, OTUB1 binds to members of the UBE2D (UBCH5) and UBE2E families, as well as to UBC13 (UBE2N). Cellular roles have been identified for the interaction of OTUB1 with UBC13 and members of the UBE2D family, but not for UBE2E E2 enzymes. We report here a novel role for OTUB1-E2 interactions in modulating E2 protein ubiquitination. We find that depletion of OTUB1 dramatically destabilizes the E2 conjugating enzyme UBE2E1 (UBE2E1) in cells and that this effect is independent of the catalytic activity of OTUB1 but depends on the ability of OTUB1 to bind to UBE2E1. We show that OTUB1 suppresses UBE2E1 autoubiquitination *in vitro* and in cells, thereby preventing UBE2E1 from being targeted to the proteasome for degradation. Taken together, we have found a new role for OTUB1 in rescuing specific E2s from degradation *in vivo*.

## Introduction

Ubiquitin signaling plays an essential role in nearly all aspects of eukaryotic biology. Substrate ubiquitination is tightly regulated temporally and in response to different stimuli through the opposing actions of enzymes that attach and remove ubiquitin from substrate proteins. Ubiquitin is conjugated to lysines via E1 ubiquitin activating enzyme, E2 ubiquitin conjugating enzymes, and E3 ubiquitin ligases, resulting in an isopeptide linkage between the ubiquitin C-terminus and a substrate lysine or, in some cases, with the substrate N-terminus (1,2). Polyubiquitin chains can be homopolymers that are linked through one of ubiquitin’s seven lysines or its N-terminal methionine, or heterotypic chains that contain a mixture of linkages (1,3). The ubiquitin modification is reversed by deubiquitinating enzymes (DUBs), which remove ubiquitin from proteins and cleave polyubiquitin chains (4,5), thus terminating a ubiquitin signaling event and recycling ubiquitin monomers. The ~90 DUBs in human cells fall into six classes based on catalytic domain architecture: ubiquitin C-terminal hydrolases (UCHs), ubiquitin-specific proteases (USP), ovarian tumor domain containing proteases (OTU), Machado Josephin domain (MJD) proteases, JAB1/MPN/Mov34 metalloproteases (JAMM), and the recently discovered MIU-containing DUB family, MINDY (4,6).

OTUB1 is an ovarian tumor domain (OTU) class cysteine protease (7) that specifically cleaves K48-linked polyubiquitin chains (8,9) and is one of the most abundant human DUBs (10). OTUB1 has been shown to regulate a diverse set of cellular processes through its deubiquitinating activity. Among the proteins that OTUB1 stabilizes are the transcription factors, FOXM1 (11) and ERα (12), and the small GTPase, RhoA (13).

OTUB1 also deubiquitinates some E3 ubiquitin ligases, such as c-IAP1, a regulator of NF-κB and MAPK signaling pathways (14), TRAF3 and TRAF6, regulators of virus-triggered interferon induction (15), and GRAIL, which regulates T-cell anergy (16).

In addition to its deubiquitinating activity, OTUB1 has the unique ability to bind to a subset of E2 ubiquitin conjugating enzymes and inhibit ubiquitin transfer in a manner that does not depend upon the catalytic activity of OTUB1 (17-19). During the double strand break response, OTUB1 binds to UBC13 and suppresses synthesis of K63-linked polyubiquitin at DNA double strand breaks, thereby regulating the DNA damage response (17). OTUB1 inhibits UBC13 by binding to the charged E2~Ub thioester intermediate and preventing ubiquitin transfer (17,19,20). OTUB1 has also been shown to non-catalytically inhibit UBCH5 in vitro (18) and in cells (21). In other examples of non-catalytic inhibition, OTUB1 stabilizes p53 by inhibiting UBCH5/MDM2-mediated ubiquitination of p53 (22,23), activates RAS isoforms (24), regulates the TGFβ pathway by stabilizing the signal transducer SMAD2/3 (25), and stabilizes DEPTOR, an mTORC1 inhibitor (26). Mass spectrometry studies have revealed that OTUB1 can form complexes with several other E2s in cells, including UBE2E1 (UBCH6), UBE2E2 (UBCH8), and UBE2E3 (UBCH9), in addition to UBC13 (UBE2N) and the UBCH5 (UBE2D1, 2, and 3) isoforms (17,27). When the E2 partners of OTUB1 are not charged with ubiquitin, E2 binding to OTUB1 stimulates its K48-specific deubiquitinating activity (28), although the physiological role of this stimulation remains to be shown.

UBE2E1 has been identified as a binding partner of OTUB1 by mass spectrometry analysis of OTUB1 binding partners in cells *(17)* and because it stimulates the deubiquitinating activity of OTUB1 *in vitro* (28). UBE2E1 is a class III E2 ubiquitin conjugating enzyme that belongs to the UBE2E family of E2s, comprising UBE2E1/UBCH6, UBE2E2/UBCH8, and UBE2E3/UBCH9 (29). UBE2E family members share a highly conserved UBC domain but are distinguished from one another by their unique N-terminal extensions. These N-terminal extensions are sites for intramolecular autoubiquitination *in vitro*, which has been shown to limit the catalytic activity of UBE2E family E2s (30,31). When lysine residues within their N-termini are mutated to arginine, or their N-termini are deleted entirely, UBE2E E2s switch from primarily monoubiquitination of substrates to robust polyubiquitination (30,31). There are several reports on the cellular functions and substrates of UBE2E1. In cells, UBE2E family E2s are imported into the nucleus when charged with ubiquitin (32). In addition to its ubiquitin conjugating activity, UBE2E1 can also act as an ISG15 conjugating enzyme *in vitro*, although this activity has not been shown to occur *in vivo* (33). UBE2E1 can be covalently modified with either ISG15 or ubiquitin, both of which interfere with the ubiquitin conjugating activity of UBE2E1 (30,33). UBE2E1 has also been reported to monoubiquitinate histone H2A at K119 in concert with the PRC1 E3 ligase complex (34).

We report here a novel role for OTUB1 in maintaining E2 levels in cells. We find that OTUB1 deficient (OTUB1^−/−^) mice exhibit late embryonic lethality. Proteomic analysis of OTUB1^−/−^ embryonic fibroblasts (MEFs) shows that levels of the E2, UBE2E1, are dramatically lower in the absence of OTUB1. In U2OS cells, knockdown or knockout of OTUB1 similarly leads to dramatically lower levels of UBE2E1 protein. This regulation of UBE2E1 stability depends on the ability of OTUB1 to bind to UBE2E1 but does not depend upon OTUB1 catalytic activity. We show that UBE2E1 is ubiquitinated *in vivo* and that in the absence of OTUB1, UBE2E1 is targeted to the proteasome for degradation. *In vitro*, UBE2E1 is autoubiquitinated in both the presence and absence of an E3 ligase, but this autoubiquitination activity is suppressed by OTUB1. Taken together, our data suggest that OTUB1 protects UBE2E1 from degradation by inhibiting E2 autoubiquitination activity. These observations reveal a novel role for OTUB1 binding to E2 ubiquitin conjugating enzymes in regulating E2 stability within the cell.

## Results

### Deletion of OTUB1 causes late stage embryonic lethality in mice

OTUB1 is the most abundant DUB in mouse and human cells and is present at concentrations of about 1 µM (10). To gain insight into the function of OTUB1 *in vivo*, we generated *OTUB1^+/−^* mice by gene targeting in C57BL/6 embryonic stem cells (Supplementary Figures 1A & 1B). Heterozygous *OTUB1^+/−^* mice were then interbred to study the phenotypic effects of *OTUB1* deficiency in vivo. We screened 124 live born mice and found that 35% were OTUB1^+/+^, 65% were heterozygous *OTUB1^+/−^* and 0% were homozygous *OTUB1^−/−^* mice (Table 1). As we found no live born OTUB1^−/−^ mice, we sacrificed pregnant females from timed matings of *OTUB1^+/−^* mice at day 14.5 of gestation, and genotyped the E14.5 embryos. These analyses revealed that the mutant *OTUB1^−^ allele* segregated in perfect Mendelian ratio (1:2:1) among 110 embryos (Table 1). These results indicate that *OTUB1* deficiency causes lethality in the late stages of embryonic development. We then used E14.5 embryos to generate *OTUB1*^−/−^ and control MEFs.

**Figure 1:**
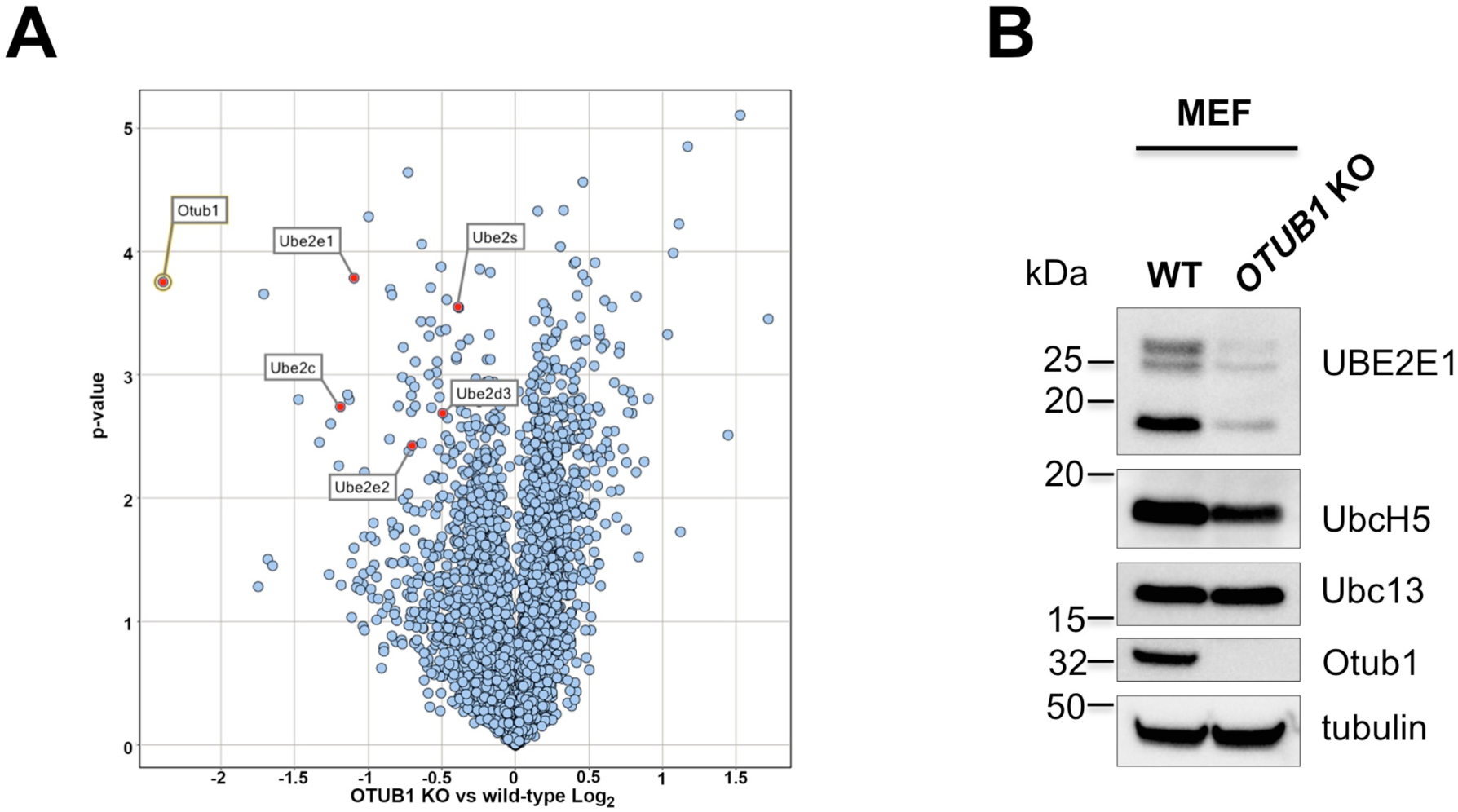
UBE2E1 levels are reduced in *OTUB1* knockout Mouse Embryonic Fibroblasts (MEFs). (A) Tandem mass tag mass spectrometry analysis of MEF wild type and *OTUB1*-/- knockout cells. (B) Western blot of the whole cell lysate of MEF WT and *OTUB1*-/- knockout cells with indicated antibodies.

**Table 1:** Embryonic Lethality of OTUB1 Deficient Mice. Numbers of embryos or live born mice of the indicated genotypes obtained from OTUB1+/- interbreeding. Percentages of each genotype indicated within parentheses. Total numbers of pups indicated at right.

|  | **OTUB1<sup>+/+</sup>** | **OTUB1<sup>+/-</sup>** | **OTUB1<sup>-/-</sup>** | **Total** |
| --- | --- | --- | --- | --- |
| **E14.5** | 29 (26%) | 51 (46%) | 30 (27%) | 110 (100%) |
| **Live born** | 44 (35%) | 80 (65%) | 0 (0%) | 124 (100%) |

### Depletion of OTUB1 destabilizes UBE2E1

We took advantage of the ability to generate *OTUB1^−/−^* null mouse embryonic fibroblasts (MEFs) in order to search for proteins whose stability depends on the presence of OTUB1. Thus far, there are only a handful of proteins whose stability is known to be regulated by OTUB1, the majority by a non-catalytic mechanism (21,25,26). We used tandem mass tag mass spectrometry (35) to search for proteins whose stability was altered in the absence of OTUB1. Proteins were extracted from both wild-type and *OTUB1*^−/−^ knockout mouse embryonic fibroblasts (MEF) and then modified with tandem mass tags prior to digestion with trypsin and liquid chromatography/mass spectrometry (LC/MS) analysis. A volcano plot of the relative changes in protein abundance in knockout versus wild type cells (Figure 1A) revealed that levels of several E2 ubiquitin conjugating enzymes are lower in *OTUB1* knockout cells, including UBE2E1/UBCH6, UBE2E2/UBCH8, UBE2C/UBCH10, UBE2S, and UBE2D3/UBCH5C. The most significant effect was on UBE2E1, which is known to bind to OTUB1 in cells (17,27).

To confirm our results, we used immunoblotting to compare steady state levels of UBE2E1 in whole cell lysates of wild-type and *OTUB1^−/−^* knockout MEFs (Figure 1B). Consistent with the mass spectrometry results (Figure 1A), we observed markedly lower UBE2E1 protein levels in *OTUB1^−/−^* MEFs as compared to wild-type cells (Figure 1B). By contrast, the OTUB1 knockout has a minimal effect on the levels of UBCH5 (UBE2D; all isoforms) and no effect on UBC13 (UBE2N) (Figure 1B), two known binding partners of OTUB1 in cells (17,27).

To ascertain whether the effect on UBE2E1 levels of an OTUB1 deletion was specific to MEFs, we used CRISPR-Cas9 (36,37) to knock out *OTUB1* gene expression in human osteosarcoma (U2OS) cells. As shown in Figure 2A, UBE2E1 levels were dramatically reduced in the U2OS knockout. To verify that the reduced UBE2E1 levels were not an adaptation to the absence of OTUB1, we used siRNA to transiently knock down OTUB1 expression in U2OS cells. As shown in Figure 2B, siRNA knockdown of OTUB1 expression similarly reduced levels of UBE2E1 (Figure 2B). Taken together, these results suggest that OTUB1 regulates UBE2E1 stability and that this regulation is not specific to a particular cell type.

**Figure 2:**
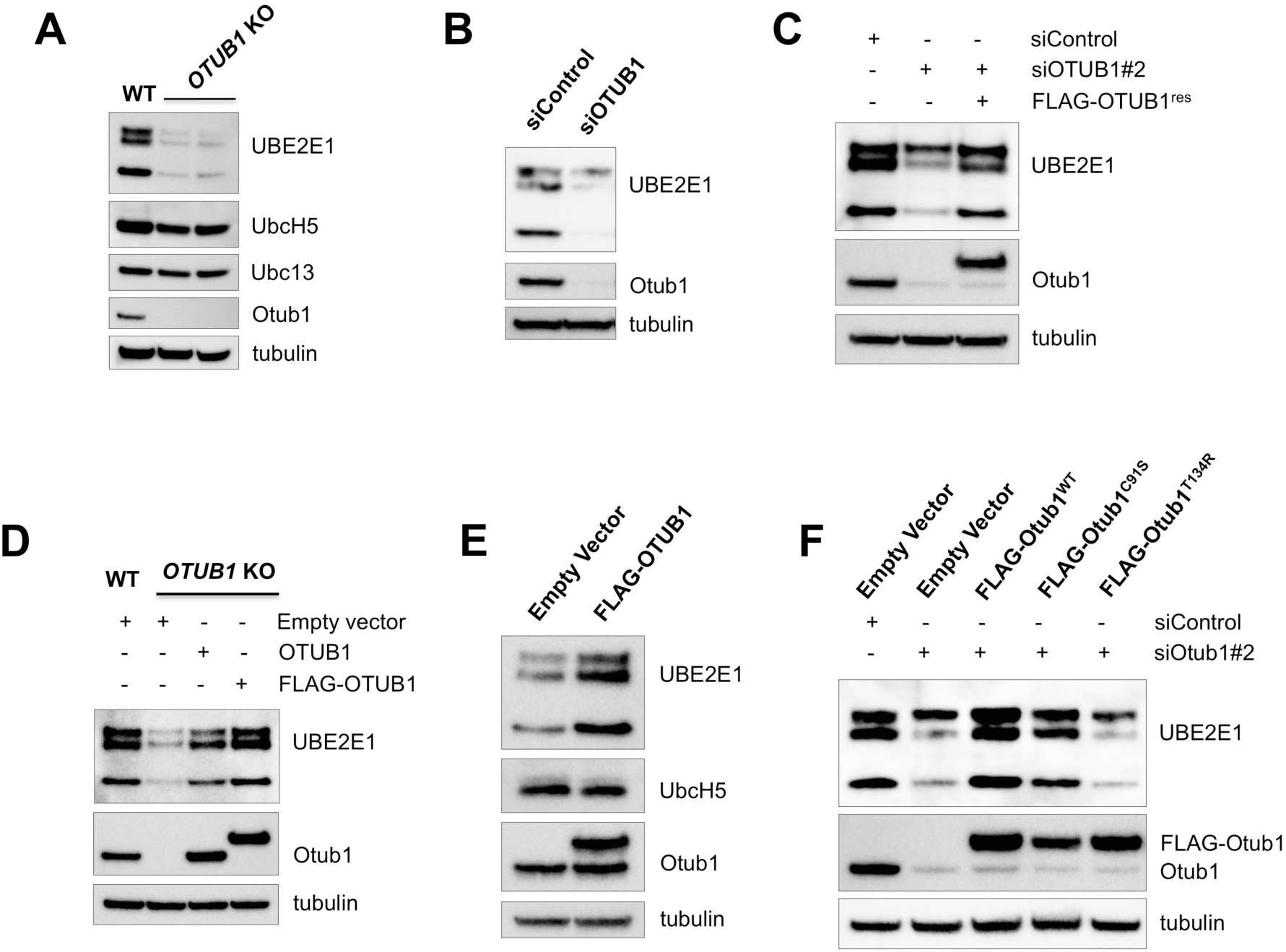
OTUB1 regulates the stability of UBE2E1 protein levels. (A) Western blot analysis of the whole cell lysate of U2OS wild-type (WT) and CRISPR-Cas9 based *OTUB1* knockout (KO) cells. (B) U2OS cells were transfected with control or SMART pool siRNA against the *OTUB1* gene. Whole cell lysates were then analyzed for the expression of indicated proteins by immunoblotting. (C) Expression of siRNA resistant FLAG-OTUB1 rescues UBE2E1 protein levels in *OTUB1* knockdown cells. U2OS stable cell lines with control plasmid or FLAG-OTUB1^res^ were transfected with non-target siRNA or individual siRNA against endogenous *OTUB1* gene and assayed for the steady-state levels of indicated proteins by immunoblotting. (D) Introduction of untagged OTUB1 or FLAG-OTUB1 in *OTUB1* knockout U2OS cells rescues UBE2E1 levels. (E) Over-expression of OTUB1 enhances UBE2E1 proteins levels. U2OS WT cells were transiently transfected with either empty vector or FLAG-OTUB1 plasmid and whole cell lysate was analyzed by western hybridization. (F) Wild-type FLAG-OTUB1^WT^ and catalytic mutant FLAG-OTUB1^C91S^ but not E2 interacting mutant FLAG-OTUB1^T134R^ rescued the steady state levels of UBE2E1 in OTUB1 knockdown U2OS cells.

We note that the anti-UBE2E1 antibody (see Methods) reacts with three different molecular weight bands in Western blots (Figure 1), irrespective of cell line, as has been observed in previous studies (34,38,39). Whereas the canonical UBE2E1 sequence has a predicted molecular weight of 21.4 kDa, the *UBE2E1* gene can express three predicted isoforms via alternative splicing. To see whether the three cross-reacting bands constituted isoforms of UBE2E1, we tested whether these proteins could be charged with ubiquitin in cells. We extracted whole cell lysates under conditions that preserve the thioester linkages (28,40) and analyzed the results by gel electrophoresis.

All three proteins that cross-react with the anti-UBE2E1 antibody migrate as higher molecular weight species in the absence of reducing agent (Supplementary Figure S2A), consistent with E2 ubiquitin conjugating activity. When compared with the molecular weight of recombinant UBE2E1 (Supplementary Figure S2B), the middle band corresponds to the canonical UBE2E1 protein while the other two bands are likely products of alternative splicing of the *UBE2E1* gene or the other UBE2E family members, UBE2E2 and UBE2E3.

### Expression of wild type or catalytically inactive OTUB1 restores UBE2E1 levels

To confirm that OTUB1 directly regulates the stability of UBE2E1, we generated stable cell lines expressing siRNA-resistant FLAG-OTUB1 and knocked down endogenous OTUB1 by siRNA. The expression of siRNA resistant FLAG-OTUB1 restores UBE2E1 protein levels when OTUB1 is knocked down (Figure 2C). UBE2E1 protein levels could also be restored in OTUB1 CRISPR knockout cell lines by expressing either FLAG-OTUB1 or untagged OTUB1 (Figure 2D). To further test whether OTUB1 stabilizes UBE2E1, we overexpressed FLAG-OTUB1 in U2OS cells and probed for UBE2E1. As shown in Figure 2E, overexpression of OTUB1 significantly enhanced the steady-state levels of UBE2E1 (Figure 2E). By contrast, levels of UBCH5 were unaffected when OTUB1 was overexpressed (Figure 2E). Taken together, our results confirm that OTUB1 specifically regulates the stability of the E2 ubiquitin conjugating enzyme, UBE2E1.

To test whether the catalytic activity of OTUB1 is required for its regulation of UBE2E1 levels, we assayed the ability of OTUB1 mutants defective in non-catalytic inhibition to complement siRNA knockdown of endogenous OTUB1 expression. Substitution of the active site cysteine with serine (C91S) inactivates OTUB1 (9) while a T134R substitution abrogates OTUB1 binding to E2 ubiquitin conjugating enzymes (18,28), thereby disrupting the ability of OTUB1 to prevent ubiquitin transfer. As shown in Figure 2F, expression of a catalytic mutant, FLAG-OTUB1^C91S^, restores levels of UBE2E1, whereas expression OTUB1^T134R^, fails to restore wild type levels of UBE2E1 (Figure 2F). We therefore conclude that OTUB1 regulates UBE2E1 levels through the previously reported ability of OTUB1 to bind to E2 enzymes (18–20), and not as a consequence of OTUB1 DUB activity.

### OTUB1 rescues UBE2E1 from proteasomal degradation

Since previous studies had identified a role for OTUB1 in stabilizing the transcription factor FOXM1 in cells (41), we considered the possibility that OTUB1 may regulate UBE2E1 protein levels at the level of transcription. We extracted total mRNA from wild-type U2OS cells, a CRISPR OTUB1 knockout, and siRNA OTUB1 knockdown cells and compared transcript levels for OTUB1 and UBE2E1 to that of TATA-box-binding protein (TBP). UBE2E1 transcript levels are not affected by OTUB1 knockout or knockdown (Supplementary Figure S3), indicating that OTUB1 does not regulate UBE2E1 levels at the level of transcription.

**Figure 3:**
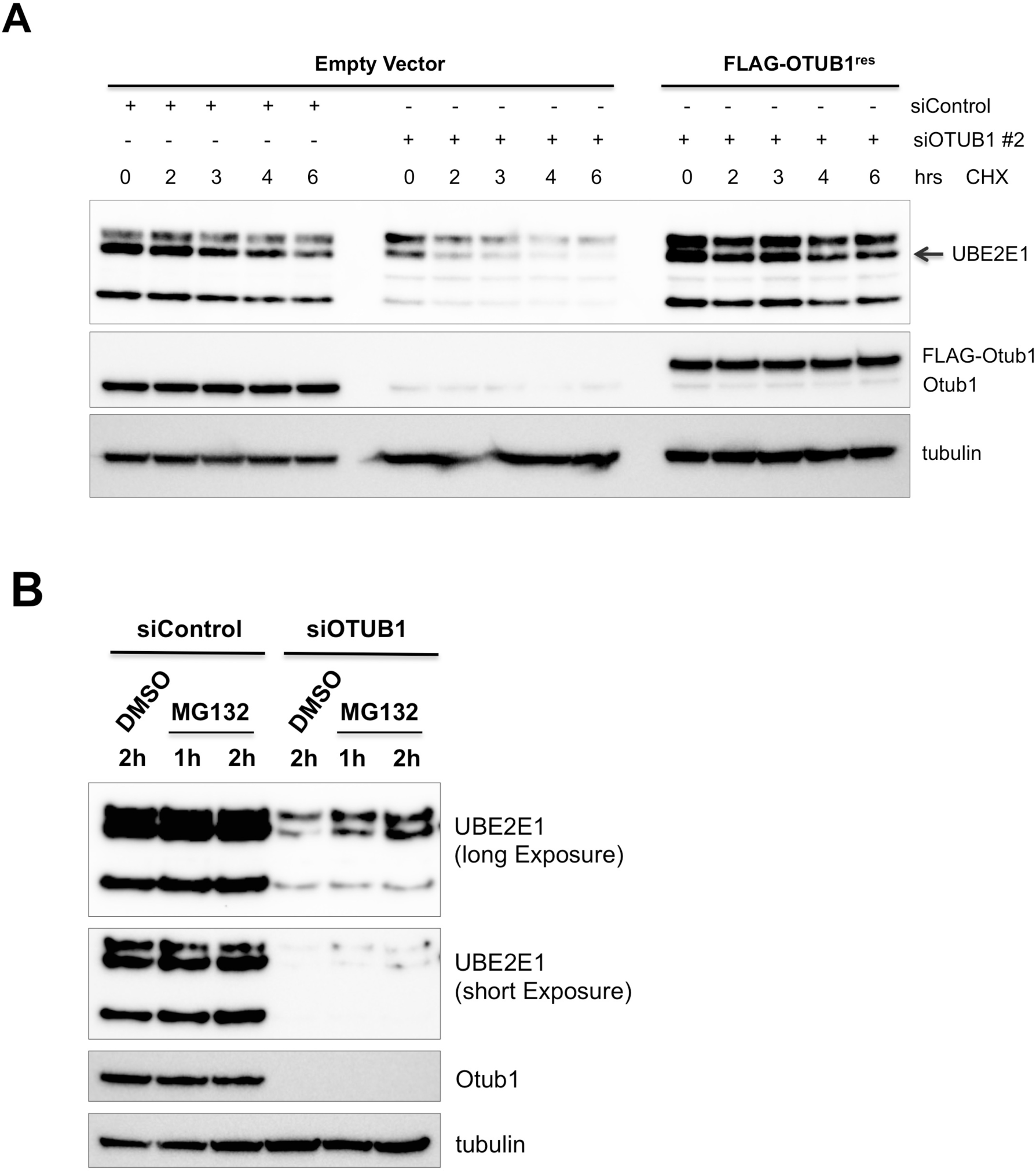
OTUB1 rescues UBE2E1 from proteasomal degradation. (A) Cycloheximide chase to assess the half-life of UBE2E1 in control and OTUB1 knockdown U2OS cells. Expression of siRNA resistant OTUB1 increases the half-life of UBE2E1 in OTUB1 knockdown cells. (B) Control and *OTUB1* knockdown U2OS cells were treated with MG132 (10µM) for 1 and 2 hr time points and whole cell lysate was assayed for the steady state levels of UBE2E1 by immunoblotting.

To test more directly whether OTUB1 stabilizes the UBE2E1 protein, we treated cells with cycloheximide to inhibit protein synthesis and monitored UBE2E1 stability in siRNA knockdown versus control cells. As shown in Figure 3A, UBE2E1 has a shorter half-life when endogenous OTUB1 expression is knocked down. Expression of FLAG-OTUB1 stabilized UBE2E1 as compared to cells in which OTUB1 is depleted. To test whether UBE2E1 is degraded by the proteasome, we treated cells with the proteasome inhibitor, MG132, and compared steady state levels of UBE2E1 in cells in which OTUB1 expression was knocked down by siRNA as compared to cells transfected with siRNA control. As shown in Figure 3B, UBE2E1 levels increase in the OTUB1 knockdown cells after two hours of MG132 treatment as compared to cells treated with DMSO. Taken together, our results suggest that OTUB1 protects UBE2E1 from proteasomal degradation through the ability of OTUB1 to non-catalytically inhibit E2 enzymes (18–20).

### OTUB1 inhibits UBE2E1 autoubiquitination

In previous studies in which OTUB1 has been shown to stabilize a substrate through its non-catalytic activity, OTUB1 inhibits the activity of an E2 that conjugates K48-linked polyubiquitin to the substrate (21–26). Since UBE2E1 is known to bind OTUB1 (17), this raises the possibility that OTUB1 may stabilize UBE2E1 directly by forming an OTUB1-UBE2E1 complex. While it has been demonstrated that UBE2E1 can stimulate OTUB1 DUB activity (28), it has not been shown whether OTUB1 inhibits UBE2E1 in the manner observed for UBC13, UBE2D2 and UBE2D3 (17,18,20). Recently, UBE2E1 was shown to autoubiquitinate itself *in vitro* in the presence or absence of the E3 ligase, RNF4 (30,31), raising the possibility that OTUB1 may stabilize UBE2E1 by preventing autoubiquitination.

To test the hypothesis that OTUB1 may directly inhibit UBE2E1, we assayed the autoubiquitinating activity of UBE2E1 in the presence and absence of OTUB1 *in vitro*. In the absence of an E3 ligase, UBE2E1 primarily monoubiquitinates itself (Figure 4, top panel). Addition of the E3 ligase RNF4 stimulates synthesis of polyubiquitin chains by UBE2E1 (Figure 4, top and middle panel). Immunoblotting with an antibody specific for UBE2E1 shows smears characteristic of polyubiquitin chains attached to UBE2E1, while immunoblotting with an antibody specific for K48-linked polyubiquitin shows the presence of higher molecular weight K48-linked polyubiquitin (Figure 4 bottom panel). The K48 chains may either be anchored to UBE2E1 or RNF4, although we cannot rule out the possibility that RNF4 stimulates UBE2E1 to form free K48-linked chains. Catalytically inactive OTUB1 (C91S) completely inhibits UBE2E1 ubiquitination activity in both the presence and absence of E3 (Figure 4 top panel). These results confirm that OTUB1 can non-catalytically inhibit UBE2E1 as has been observed for UBC13 and UBCH5 isoforms (17–20).

**Figure 4.**
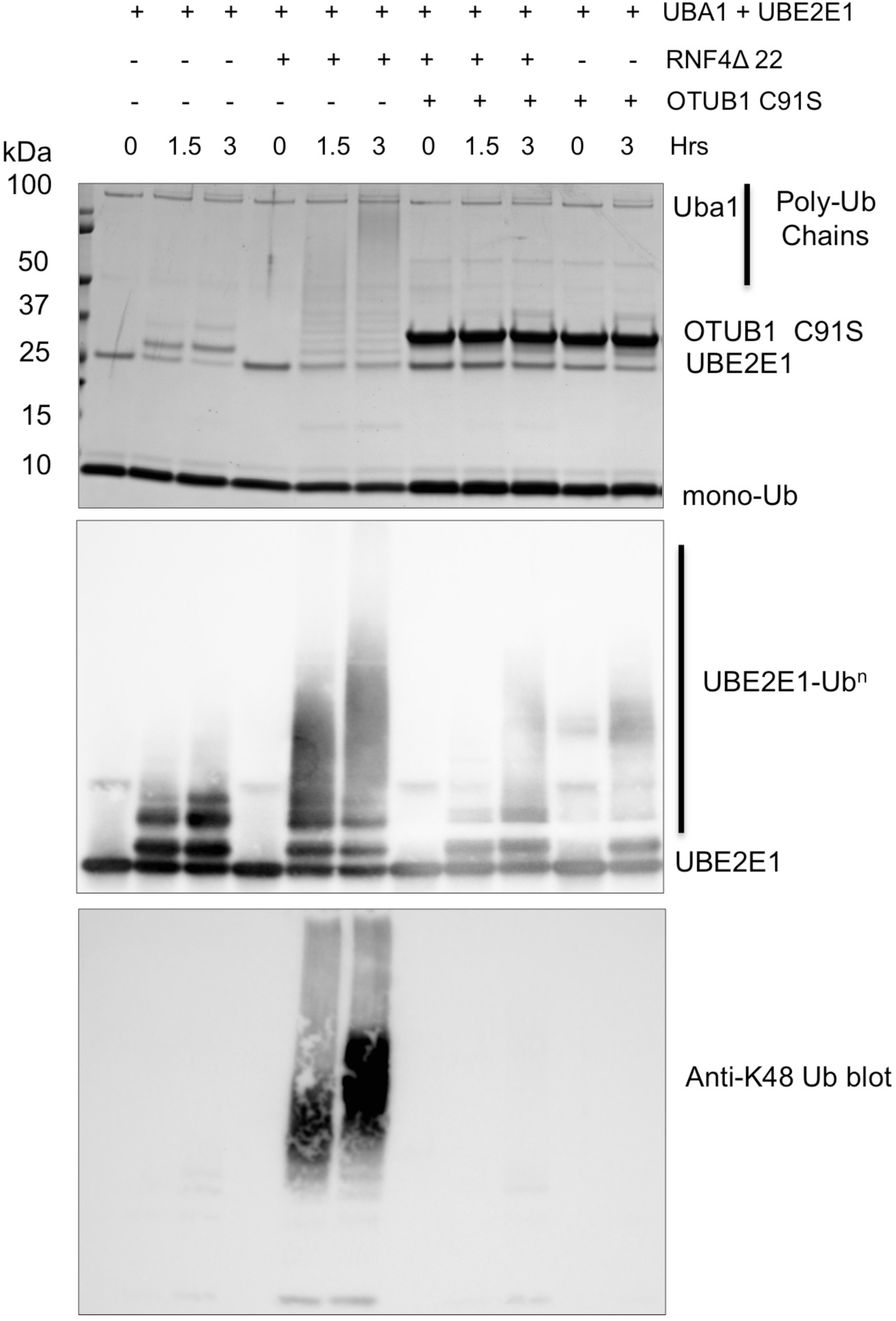
OTUB1 suppresses autoubiquitination of UBE2E1. (A) In vitro assay of recombinant proteins showing autoubiquitination of UBE2E1 in the presence and absence of the E3 ligase, RNF4, and OTUB1-C91S. Coomassie stained gel of ubiquitination reactions containing 0.1 nM UBA1, 5 µM UBE2E1, 1 µM RNF4 Δ22, 50 µM Ubiquitin, and 10 µM OTUB1 C91S. (B) Western blot of reactions shown in panel (A) using antibodies against UBE2E1 (middle) and K48-polyubiquitin (bottom).

While it is not known whether RNF4 stimulates UBE2E1 *in vivo*, our *in vitro* results raised the interesting possibility that OTUB1 could inhibit UBE2E1 autoubiquitination in cells, thereby accounting for the ability of OTUB1 to prevent UBE2E1 from begin targeted to the proteasome. To test this hypothesis, we co-transfected U2OS cells with plasmids expressing HA-tagged UBE2E1 and His_6_-tagged ubiquitin, and then treated the cells with MG132 to enrich for ubiquitinated proteins. His_6_-tagged ubiquitinated proteins were pulled down by Ni^2+^ NTA resin under denaturing conditions and the results analyzed by Western blots were probed with an anti-HA antibody. We found that HA-UBE2E1 is primarily monoubiquitinated in cells, with a small amount of higher molecular weight bands also observed (Figure 5A). To determine whether monoubiquitination of UBE2E1 in cells is due to autoubiquitination activity, we compared autoubiquitination of wild-type HA-UBE2E1 and a catalytically inactive mutant HA-UBE2E1^C131A^ under the same experimental conditions as described above. As shown in Figure 5B, wild-type HA-UBE2E1 is ubiquitinated but not the catalytically inactive mutant HA-UBE2E1^C131A^, supporting the idea that UBE2E1 is autoubiquitinated in cells.

**Figure 5:**
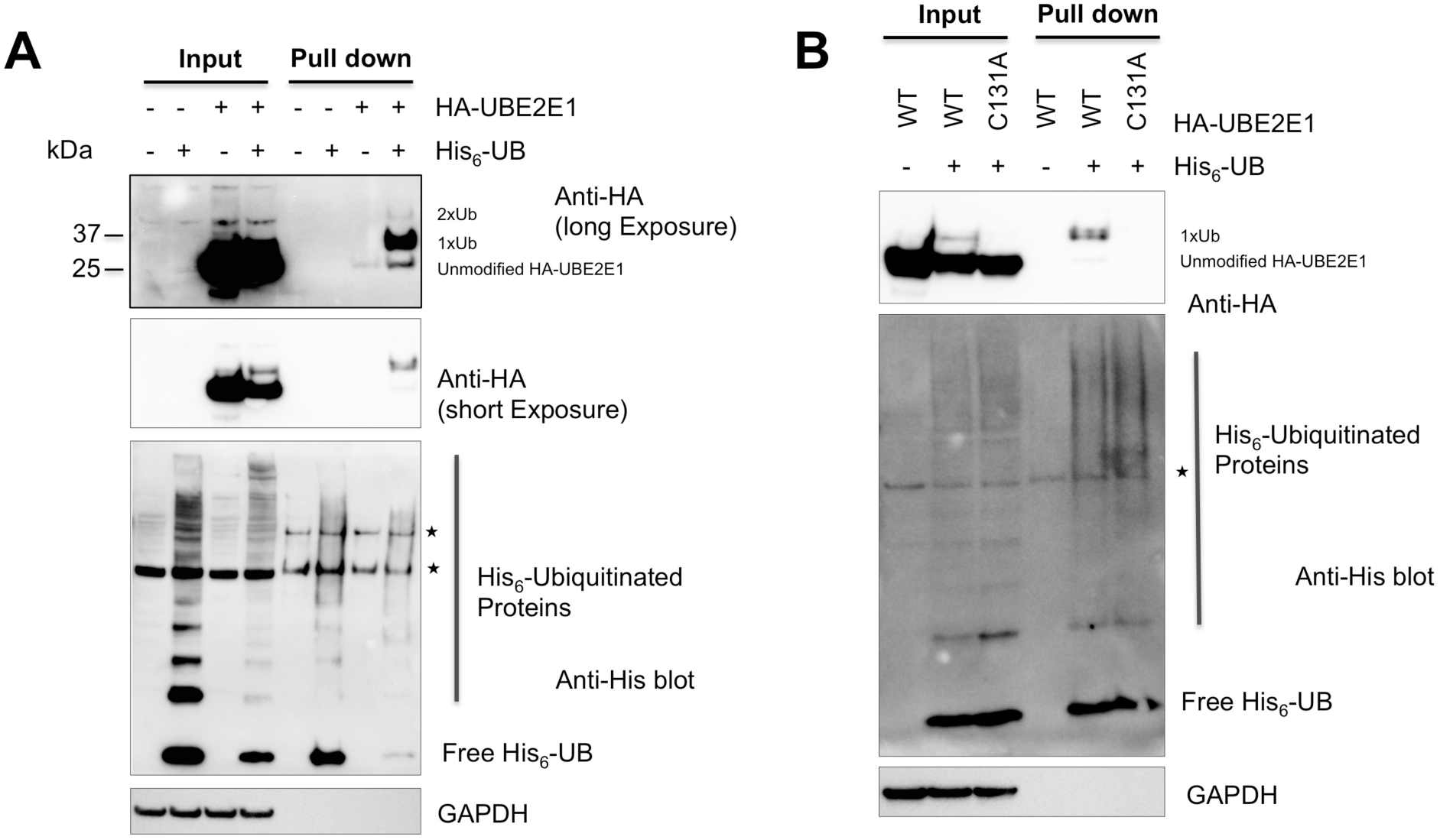
UBE2E1 is auto-ubiquitinated in U2OS cells. (A) U2OS cells were co-transfected with plasmids expressing HA-UBE2E1 and His_6_-Ubiquitin. His_6_ tagged ubiquitinated proteins in the whole cell lysate were enriched using Ni^2+^ NTA agarose beads and analyzed by Western hybridization with anti-HA antibody. (B) U2OS cells were co-transfected with plasmids expressing either wild-type HA-UBE2E1 or catalytically inactive HA-UBE2E1 (C131A) and His_6_-Ubiquitin. His_6_ tagged ubiquitinated proteins and whole cell lysate analyzed as in (A). Asterisk denotes cross-reactive bands.

### Histone ubiquitination is not affected by OTUB1 or UBE2E1

UBE2E1 has been reported to monoubiquitinate histone H2A at K119 (34), a histone mark that regulates gene silencing (42). In light of our observation that the absence of OTUB1 destabilizes UBE2E1, we tested whether decreased cellular levels of OTUB1 reduce ubiquitination of H2A-K119. However, we find no detectable difference in levels of ubiquitinated H2A when OTUB1 is knocked down by siRNA in U2OS cells (Figure 6A and S4A), or in a CRISPR-Cas9 OTUB1 knockout cell line (Figure 6B and S4B). As a control to rule out possible indirect effects impacting H2A ubiquitination levels, we used siRNA to knock down expression of UBE2E1 directly and examined the steady state levels of ubiquitinated H2A. Contrary to a previous report (34), we did not find that UBE2E1 siRNA knockdown affected ubiquitination of H2A-K119 (Figure 6C and S4C). We similarly found that a UBE2E1 knockdown did not impact histone H2B monoubiquitination (Figure 6C), a mark of actively transcribed chromatin (43). While we cannot rule out additional factors that may account for the lack of an effect of UBE2E1 or OTUB1 knockdown on histone H2A monoubiquitination, we note that the RING1B/BMI1 E3 ligase complex that monoubiquitinates H2A-K119 also ubiquitinates H2A together with the E2, UBCH5C (44,45), which may compensate for the absence of UBE2E1 in our experiments. Our results are therefore more consistent with a role for UBCH5C, rather than UBE2E1, in monoubiquitinating H2A-K119.

**Figure 6:**
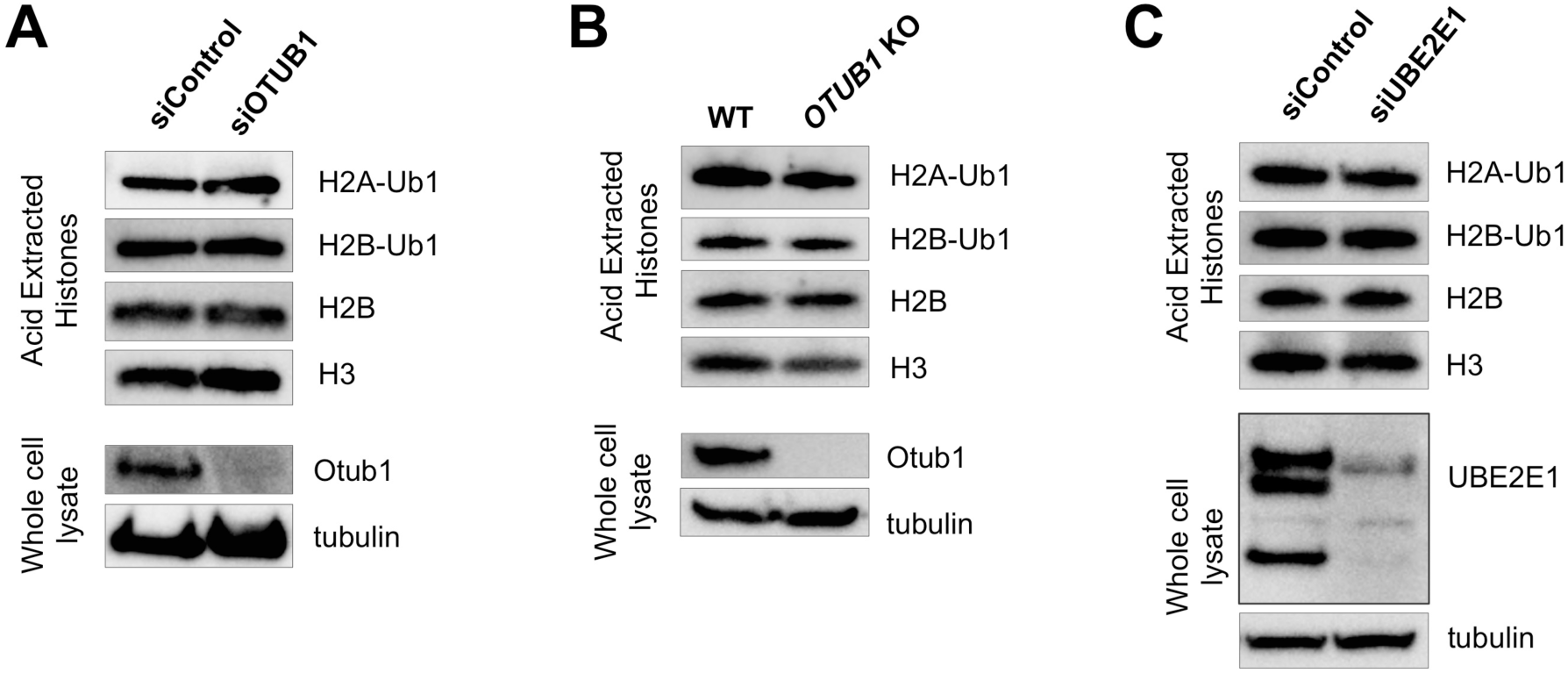
Depletion of OTUB1 or UBE2E1 does not affect global H2A-Ub and H2B-Ub levels. Whole cell lysates and acid-extracted histones were analyzed by immunoblotting with the indicated antibodies. (A) Control and OTUB1 knocked down U2OS cells. (B) U2OS WT and CRISPR OTUB1 knockout cells. (C) Control and UBE2E1 knockdown U2OS cells.

## Discussion

The ability of OTUB1 to bind to E2 enzymes and inhibit ubiquitin transfer in a manner that does not depend on OTUB1 catalytic activity was first discovered in studies of DNA damage signaling, in which OTUB1 inhibits the E2, UBE2N/UBC13 (17). OTUB1 was also shown to bind to members of the UBE2D and UBE2E E2 families, although the in vivo significance of that observation was not known (17,20,28). Subsequent studies have shown that OTUB1 stabilizes proteins such as p53 (21,22), SMAD2/3 (25), and DEPTOR (26) in a non-catalytic manner by preventing transfer of ubiquitin from an E2 to the substrate. The E2 that ubiquitinates these substrates in trans are thought to belong to the UBE2D (UBCH5) family, based on the promiscuous activity of these E2 enzymes with most E3 ligases and the ability of OTUB1 to inhibit members of this family (17,18). In this study, we show that OTUB1 stabilizes UBE2E1 (UBCH6) in a unique manner by preventing this E2 from ubiquitinating its own lysines, thereby protecting UBE2E1 from proteasomal degradation.

The ability of OTUB1 to suppress UBE2E1 autoubiquitination explains why knockdown or knockout of OTUB1 dramatically stabilizes UBE2E1 protein levels but has little to no effect on levels of other E2 partners of OTUB1 such as UBC13 and UBCH5 isoforms (17,18,20) (Figure 1). Previous studies have shown that UBE2E1 is autoubiquitinated *in vitro* in the absence or presence of an E3 ligase (30,31), although the relevance of this activity *in vivo* was not clear. We show here that OTUB1 non-catalytically inhibits UBE2E1 autoubiquitination *in vitro* in either the presence or absence of E3 ligase (Figure 4). Analysis of whole cell lysates showed that the dramatic reduction in UBE2E1 protein levels in OTUB1 knockout or knockdown cells could be rescued by expressing either wild type OTUB1 or its catalytically inactive mutant, C91S, but not the OTUB1 T134R mutant that is defective in binding to E2 enzymes (18,28) (Figure 2). When OTUB1 is depleted, autoubiquitination of UBE2E1 leads to rapid degradation by the proteasome and shortened half-life in the cell (Figure 3). In pull-downs of ubiquitinated proteins from cells, we found that wild type HA-UBE2E1 was ubiquitinated, but not the catalytically inactive mutant, HA-UBE2E1 C131A (Figure 5), consistent with autoubiquitination in cells. Taken together, these results support a role for OTUB1 in regulating autoubiquitination and proteolytic degradation of UBE2E1.

UBCH6/UBE2E1 belongs to the UBE2E family of E2s, comprised of UBE2E1, UBE2E2, and UBE2E3. Our proteomics data uncovered lower protein levels for UBE2E2 in addition to UBE2E1 (Figure 1A), indicating that OTUB1 probably also regulates the stability of this E2. Since all members of the UBE2E family can autoubiquitinate themselves *in vitro* (30,31), the mechanism of stabilization we observed for OTUB1-UBE2E1 likely extends to UBE2E2 and UBE2E3. All three UBE2E family members can interact with a broad set of E3 ubiquitin ligases (46), suggesting that OTUB1 may have further downstream effects on other substrates that are ubiquitinated by UBE2E enzymes.

We have also demonstrated that neither OTUB1 nor UBE2E1 has a detectable effect on ubiquitination of histone H2A, contrary to a recent report (34). Despite assays of multiple biological replicates in OTUB1 siRNA knockdown and CRISPR-Cas9 knockout U2OS cells, we detected no difference in histone ubiquitination using highly selective antibodies against H2A-K119Ub. We speculate that differences in methods for extracting histones may give rise to these discrepancies. In this study, we analyzed histones that were acid extracted and fully solubilized from chromatin fractions, and normalized loading relative to histone H3. Based on our observations, we hypothesize that UBE2E1 is likely not the primary cognate E2 conjugating enzyme for the PRC1 E3 ligase. The UBCH5 isoforms ubiquitinate histone H2A in concert with PRC1 *in vitro* (44,45), and TRIM37, an E3 ligase that associates with PRC2, ubiquitinates H2A in human breast cancer cell lines where RING1B of PRC1 is downregulated (47). Further experiments targeting both UBE2E1 and the UBCH5 isoforms may shed light on which E2 is primarily responsible for H2A-K119 ubiquitination in cells.

The results of our OTUB1 knockout in mice indicate that OTUB1 plays a critical role in embryonic development. However, the molecular basis for this late embryonic lethal phenotype is currently unclear. Based on our cell culture experiments, it is tempting to speculate that OTUB1 depletion causes embryonic lethality by destabilizing UBE2E1 and impairing ubiquitination of its targets, one of which might play an essential role in embryogenesis. However, UBE2E1 knockout mice are viable (IMPC database, http://www.mousephenotype.org), making it unlikely that the mechanism we describe here is the only contributing factor to the embryonic lethal phenotype. It is possible that a combination of the catalytic activity of OTUB1 and its ability to inhibit E2 conjugating enzymes contribute to its importance in development. Further studies will be needed to determine definitively the developmental pathways that are disrupted in the absence of OTUB1.

In summary, we have shown that OTUB1 non-catalytically regulates the stability and protein levels of UBE2E1 in the cell. Evidence suggests that ubiquitin conjugating enzymes, ubiquitin ligases, and deubiquitinating enzymes exist in complex *in vivo* (27,48). Our results demonstrate that these complexes may not only serve to regulate ubiquitinated substrates, but also affect the levels of ubiquitin machinery in the cell. OTUB1 was previously known to non-catalytically prevent ubiquitination of substrates by binding to E2 conjugating enzymes. Here we have shown a new role for OTUB1 non-catalytic inhibition, namely by preventing E2 autoubiquitination that leads to proteasomal degradation. Our findings add a new layer of complexity to the increasingly complex mechanism of OTUB1-regulated ubiquitin signaling in the cell.

## Materials and Methods

### Generation of OTUB1 knockout mouse and mouse embryonic fibroblasts

The gene targeting construct (Supplementary Figure S1A) was generated from a bacterial artificial chromosome (BAC) from the C57BL/6J strain containing the OTUB1 gene by recombineering, replacing exons 4-7 with a loxP-flanked neomycin-resistance gene (Neo). C57BL/6 embryonic stem cells (PRX-B6T) were transfected with this construct and successfully targeted ES cells were identified by novel Xba I fragments identified by Southern blots (Supplementary Figure S1B). Correctly targeted clones were injected into blastocysts by the UCSF Transgenic Core. Chimeras were bred to albino C57BL/6J mice and nonalbino progeny were screened for the presence of the targeted allele by Southern blot (Otub1^+/−^). Heterozygous (Otub1^+/−^) mice were interbred and plugged females were sacrificed at day 14.5 of gestation to generate mouse embryonic fibroblasts (MEFs). All animal studies were conducted in accordance with the University of California, San Francisco Institutional Animal Care and Use Committee.

### Cell lines, plasmids and antibodies

U2OS and MEF cells were grown in Dulbecco’s modified Eagle’s medium supplemented with 10% FBS. U2OS cell line was tested for authenticity by STR profile analysis and found 100% identical to the ATCC STR profile database. The following antibodies were used for immunoblotting: Anti-OTUB1 (Abcam, Cat # ab175200), Anti-UbE2E1/UBCH6 (Boston Biochem, Cat # A630), Anti-UBCH5 (Boston Biochem, Cat # A615), Anti-UBC13 (Invitrogen, Cat # 371100), Ubiquityl-Histone H2A (Lys119) (Cell Signaling Technology, Cat # 8240), Ubiquityl-Histone H2B (Lys 120) (Cell Signaling Technology, Cat # 5546), Histone H3 (Cell Signaling Technology, 4499), Histone H2B (Cell Signaling Technology, 12364), Anti-GAPDH (Cell Signaling Technology, Cat # 5174), Anti-α-Tubulin (Sigma cat # T6199), Anti-HA (Invitrogen # 32-6700), Anti-ubiquitin (P4D1), Anti-pentaHis-HRP (Qiagen). pCI-His_6_-ubiquitin plasmid was obtained from Addgene (Cat # 31815). Construction of FLAG-OTUB1 plasmids was previously described in (9,20). HA-UBE2E1 was cloned into pCDNA between HindIII and XhoI restriction sites. Catalytically inactive HA-UBE2E1 (C131A) mutant was constructed by site-directed mutagenesis PCR.

### RNA interference

All siRNAs employed in this study were purchased from GE Dharmacon. Transfections were performed using Lipofectamine RNAimax (Life Technologies) following the manufacturer’s protocol and cells were analyzed after 3 days post-transfection. The sequences of SMARTpool siRNA for OTUB1 (catalogue # M-021061-01-0005) are 5’-GACAACAUCUAUCAACAGA-3’(siRNA#1), 5’-CCGACUACCUUGUGGUCUA-3’(siRNA#2), 5’-GACGGCAACUGUUUCUAUC-3’(siRNA#3), 5’GACGGACUGUCAAGGAGUU-3’(siRNA#4), the sequence of individual siRNA for UBE2E1 is 5’-GACCAAGAGAUACGCUACA-3’. Non-target siRNA (cat # D-001210-01-05) was used as control. Complementation assays for OTUB1 were done by introducing siRNA#2 in U2OS stable cell lines expressing siRNA resistant Flag-OTUB1^res^. To generate siRNA resistant flag-OTUB1, the target sequence in OTUB1 was re-coded to CCGACTACCTCGTTGTCTA by site-directed mutagenesis.

### CRISPR/Cas9 based OTUB1 knockout

Guide RNA targeting the exon II of OTUB1 gene was designed using online MIT tool http://crispr.mit.edu. The sequence of guide RNA is *TCGGTCCTGCTGAGCCATGA.* Oligos for guide RNA were annealed and ligated into BbsI digested pSpCas9(BB)-2A-Puro vector (Addgene # 62988) as described (37,49). Integration of the guide RNA into the plasmid was confirmed by restriction digest and sequencing. Guide RNA plasmid was transfected into U2OS cells using Lipofectamine 2000 reagent following manufacturer’s protocol and incubated for 3 days. Single cells were isolated and cultured for 3-4 weeks in Puromycin (2 µg/ml) medium to generate isogenic stable cell lines. Genomic DNA was extracted from individual colonies and guide RNA targeted exon II region was PCR amplified (FWD primer: *CTAAGCCTGTCTTCCTGACCCT* and REV primer: *AGCTTCCAAAGTAGAGACAGAC*) and sequenced to screen for the presence of indel mutations. Further, all possible exonic OFF target sites that are predicted by the MIT tool were PCR amplified and sequenced. Primers sequence for OFF target screening are listed in table.

### Mass spectrometry

Three biological replicates of wild-type and *OTUB1^−/−^* MEF cells were grown to 70% confluence, then harvested and washed with ice cold PBS. Cells were lysed by incubation in PBS/SDS lysis buffer (1xPBS, 2% SDS, 1x protease inhibitors (Roche EDTA-free), 25 mM NEM, 5 mM o-phenanthroline, 1 mM EDTA, 1 mM PMSF) at 4°C rocking for 20 minutes. Lysate was briefly sonicated, then cellular debris was pelleted by centrifugation at 13,000 rpm for 10 min at 4°C. Total protein was quantified by Pierce 660 nm protein assay. 100 μg total protein was reduced and alkylated by treatment with TCEP and MMTS, followed by TCA precipitation of proteins at −20°C. Precipitated proteins were pelleted and dried, and then submitted to the JHMI mass spectrometry core facility for trypsinization, tandem mass tag labeling, and LC/MS analysis.

### Quantification of mRNA levels by Real-Time PCR

Equal numbers of wild-type or OTUB1 CRISPR knockout U2OS cells were plated, and the next day were transfected with either control siRNA or OTUB1 SMARTpool siRNA with Lipofectamine RNAimax. Two days after transfection, cells were harvested and total RNA was extracted with a GenElute kit (Sigma Aldrich), following the manufacturer’s protocol. cDNA was synthesized from 200 ng total RNA using the NEB ProtoScript First Strand cDNA synthesis kit, following the manufacturer’s protocol. RT-PCR reactions contained 1 μL of cDNA, 500 nM forward and reverse primers, and iTaq SYBR green premix (Biorad). Reactions were run on a QuantStudio RT-PCR machine (Thermo Fisher). Relative mRNA values were quantified from ΔΔC_t_ values extracted for UBE2E1 and OTUB1 genes compared to TBP using 3 experimental and 2 technical replicates. Primers used for RT-PCR were: 5’-AGATGTTATCGCCTTTGGGA-3’ and 5’-TCCAAACTCCTCTCCACCAG-3’ for *UBE2E1*; 5’-GCCATAAGGCATCATTGGAC-3’ and 5’-AACAACAGCCTGCCACCTTA-3’ for *TBP*; 5’-AACACGTTCATGGACCTGATTG-3’ and 5’-TGCTCTGGTCATTGAAGGAGG-3’ for *OTUB1* primer set 1; and 5’-CCATCATGGCTCAGCAGGA-3’ and 5’-GAGGTCCTTGATCTTCTGTTGATAGATG-3’ for *OTUB1* primer set 2.

### Pulldown of ubiquitinated UBE2E1

U2OS cells were co-transfected with plasmids expressing HA-UBE2E1 (pCDNA) and His_6_-ubiquitin for 48 hrs. Cells were then treated with 25 µM MG132 (Cayman Chemical) for 3 hrs before harvesting. Whole cell lysate was made in denaturing conditions as described in (21). His_6_-ubiquitin conjugated species were enriched by incubating with Ni^2+^ NTA agarose beads for 3 hrs on rotator and analyzed by western blotting.

### Acid extraction of whole cell histones

After 72 h of post-transfection with siRNA, U2OS cells were harvested by trypsinization and re-suspended in culture medium. Cells were pelleted by centrifugation at 1400 rpm for 5 min at room temperature. Cell pellet was washed twice with ice-cold PBS containing 2 mM PMSF. Whole cell lysate was made by adding 100 µl of RIPA buffer containing 1x protease inhibitor (Roche) and 10 mM N-ethylmaleimide (NEM) to one-third of the cell pellet. Lysate was clarified by centrifuging at 13,000 rpm for 10 min, 4°C and boiled in SDS sample buffer for 5 min. Lysate was analyzed for efficient knockdown of the protein of interest. Remaining two-third cell pellet was used for acid extraction of histones (50). Cell pellet was resuspended in 1 mL ice-cold hypotonic lysis buffer (10 mM Tris-HCl pH 8.0, 1 mM KCl, 1.5 mM MgCl_2_, 1 mM DTT, 2 mM PMSF, 10 mM NEM and 1x Protease inhibitor cocktail) and incubated for 30 min on rotator at 4**°**C. After hypotonic lysis, intact nuclei were pelleted by centrifuging in cooled table top microfuge at 10,000xg for 10 min. Nuclei pellet was resuspended in 400 µL of ice-cold 0.4N H_2_SO_4_ and incubated on rotator for 30 min at 4°C to extract histones. Sample was spun in cooled microfuge at 16,000xg for 10 min to remove nuclear debris. Trichloroacetic acid was added to the supernatant to a final concentration of 33% and then incubated on ice for 30 min to precipitate histones. Histones were pelleted by centrifuging at 16,000xg for 10 min and 4°C. Histones were washed twice with 1 mL of ice-cold acetone and air-dried for 20 min at room temperature. Histones were dissolved in 100 µL of 1x SDS sample buffer and analyzed by western hybridization.

### Western blotting

Whole cell lysate and acid extracted histones were separated on SDS-PAGE (4-12% Bis-Tris, Pre-cast gel, Criterion XT) and then transferred to PVDF membrane by Bio-rad Gel transfer system. Membranes were blocked with 4% Bio-Rad blotting grade blocker in TBST and hybridized with indicated primary antibodies overnight at 4°C. Membrane was washed three times with TBST and incubated with horseradish peroxidase (HRP) conjugated secondary antibodies. Super Signal Pico ECL (Thermo Scientific) was used for chemiluminescent detection of horseradish peroxidase conjugated antibody. Chemiluminescent signals were detected with Bio-Rad ChemiDoc Touch imaging system.

### *In vitro* ubiquitination assays

Ubiquitination assays were done at 37°C in a buffer of 50 mM HEPES pH 7.5, 250 mM NaCl, 10 mM MgCl_2_, 0.5 mM DTT and 0.005% Tween-20. All reactions contained 0.1 nM Uba1, 5 µM UBE2E1, 1 µM RNF4Δ22, and 50 µM ubiquitin. Inhibited reactions contained 10 µM of catalytically inactive OTUB1 C91S. Assays were initiated by the addition of 5 mM ATP and Uba1. Aliquots were removed at specific time points and quenched with SDS-PAGE loading buffer containing β-mercaptoethanol. Samples were run on 4-12% polyacrylamide Bis-Tris Criterion XT gels (Bio-Rad) and stained with Coomassie brilliant blue.

## Acknowledgements

Supported by grant GM109102 from the National Institute of General Medical Sciences (C.W.).

---

**Supplementary Table 1.**
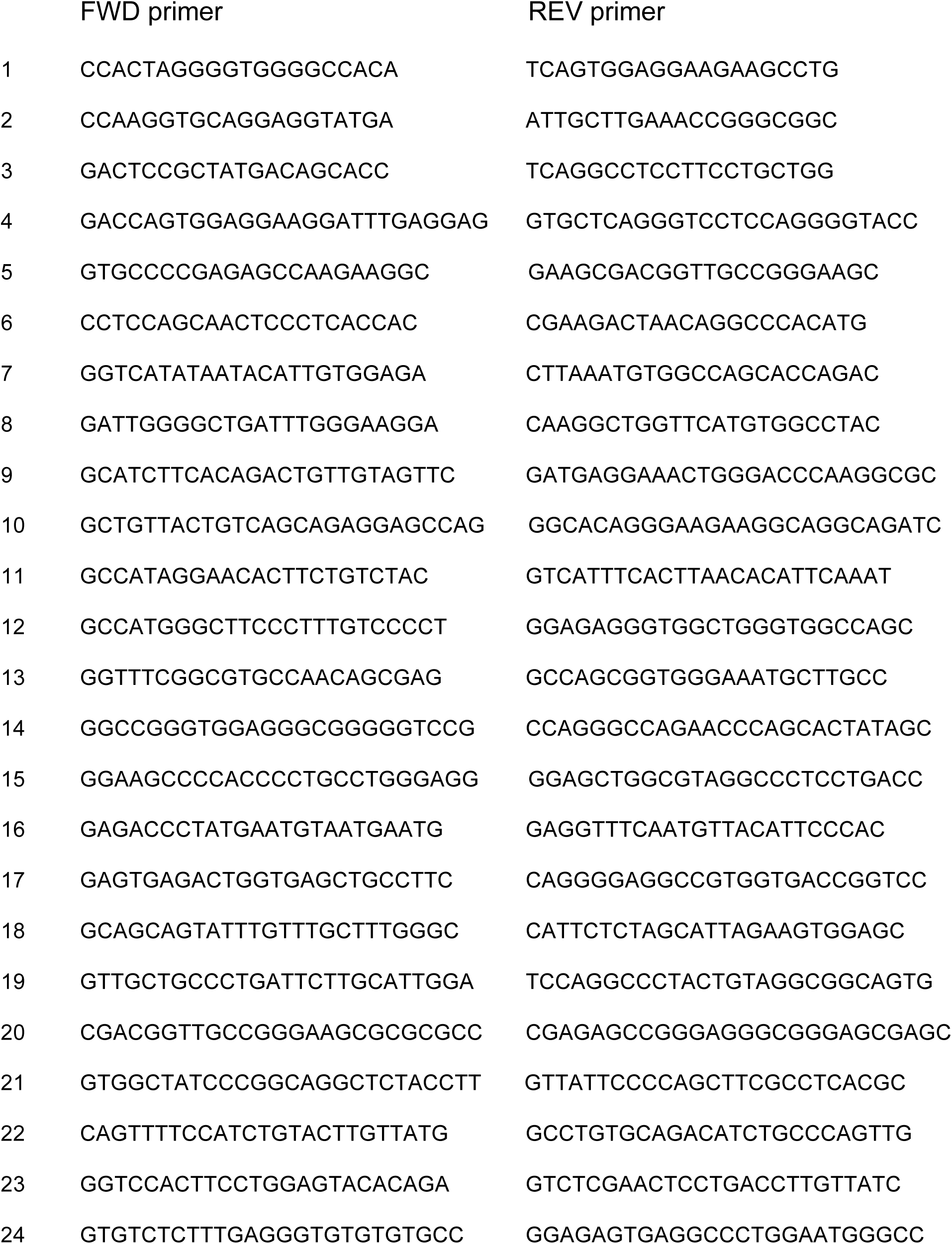
List of primers for OFF target screening

**Figure S1:**
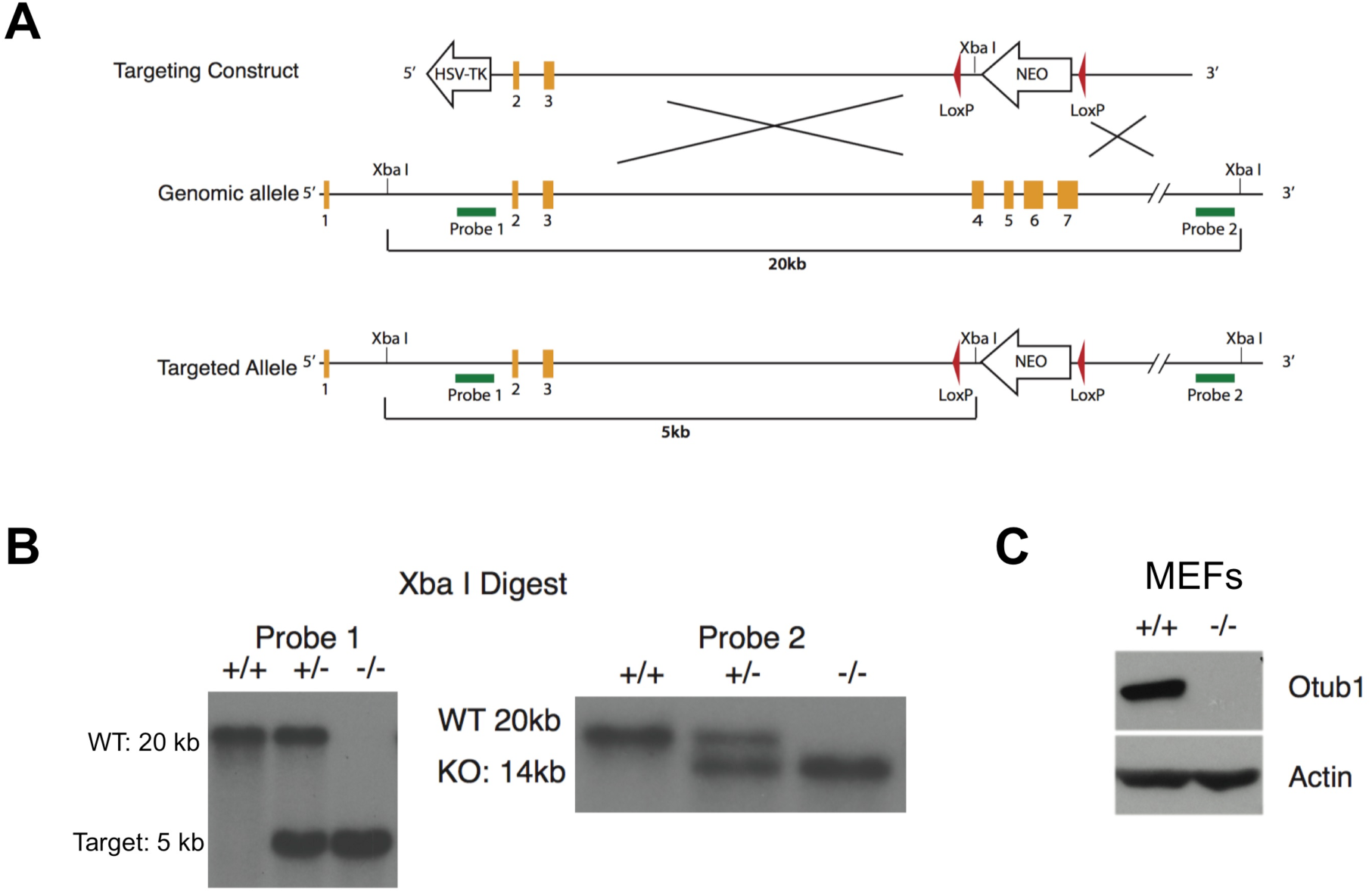
Gene targeting strategy to generate OTUB1 knockout mice. (A) Schematic representation of the gene targeting construct and screening strategy to delete exons 4-7 of OTUB1. (B) Southern blots of Xba I digested genomic DNA from targeted ES cells. (C) Immunoblot of primary MEFs cultured from OTUB1^+/+^ and OTUB1^−/−^.

**Figure S2:**
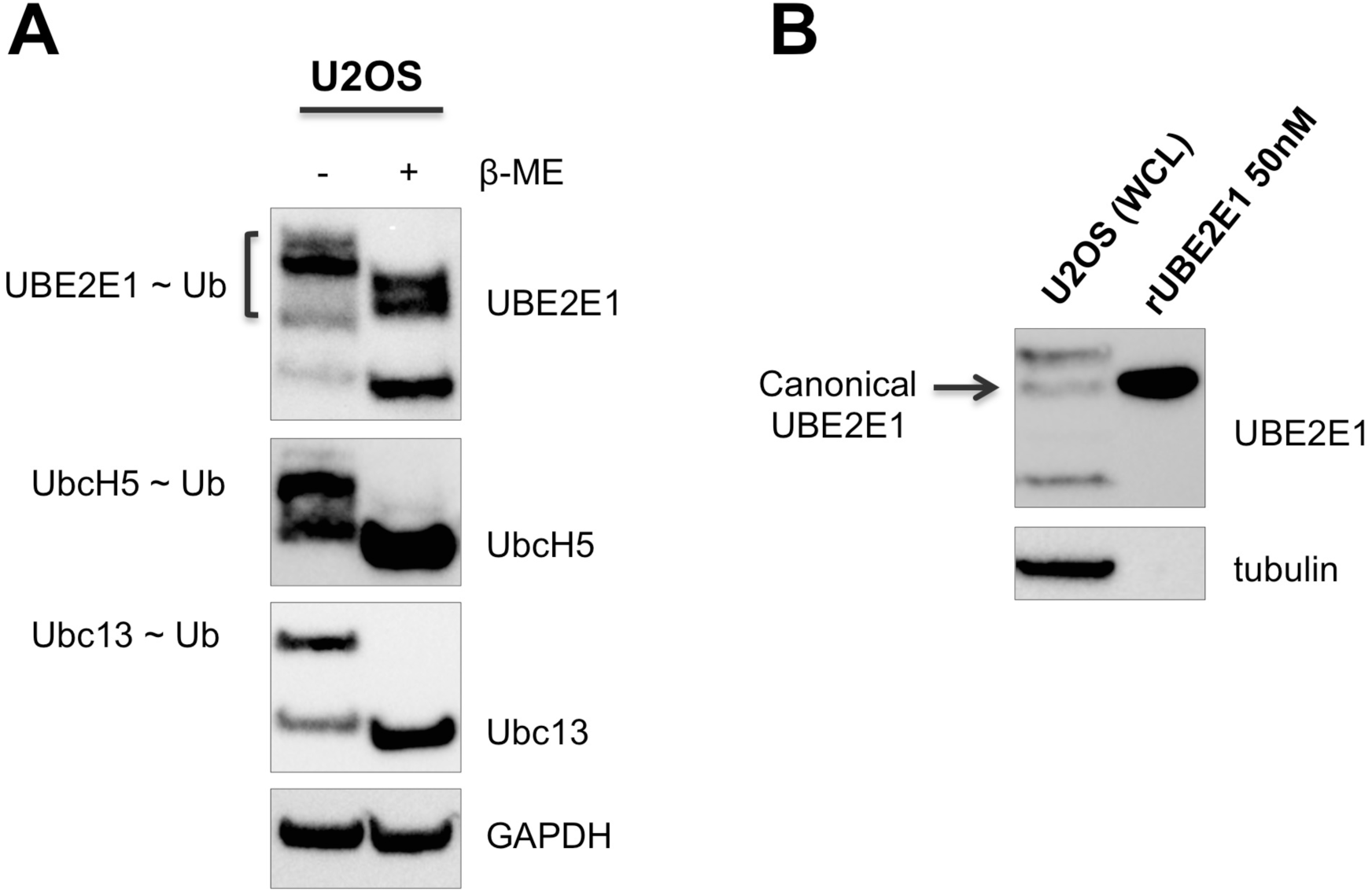
All proteins that cross-react with the anti-UBE2E1 antibody are charged with ubiquitin. (A) Whole cell extracts of U2OS cells were incubated with and without β-mercaptoethanol and analyzed by western hybridization with the indicated antibodies. (B) Whole cell extract from U2OS cells and 50 nM of recombinant UBE2E1 were analyzed by western hybridization.

**Figure S3:**
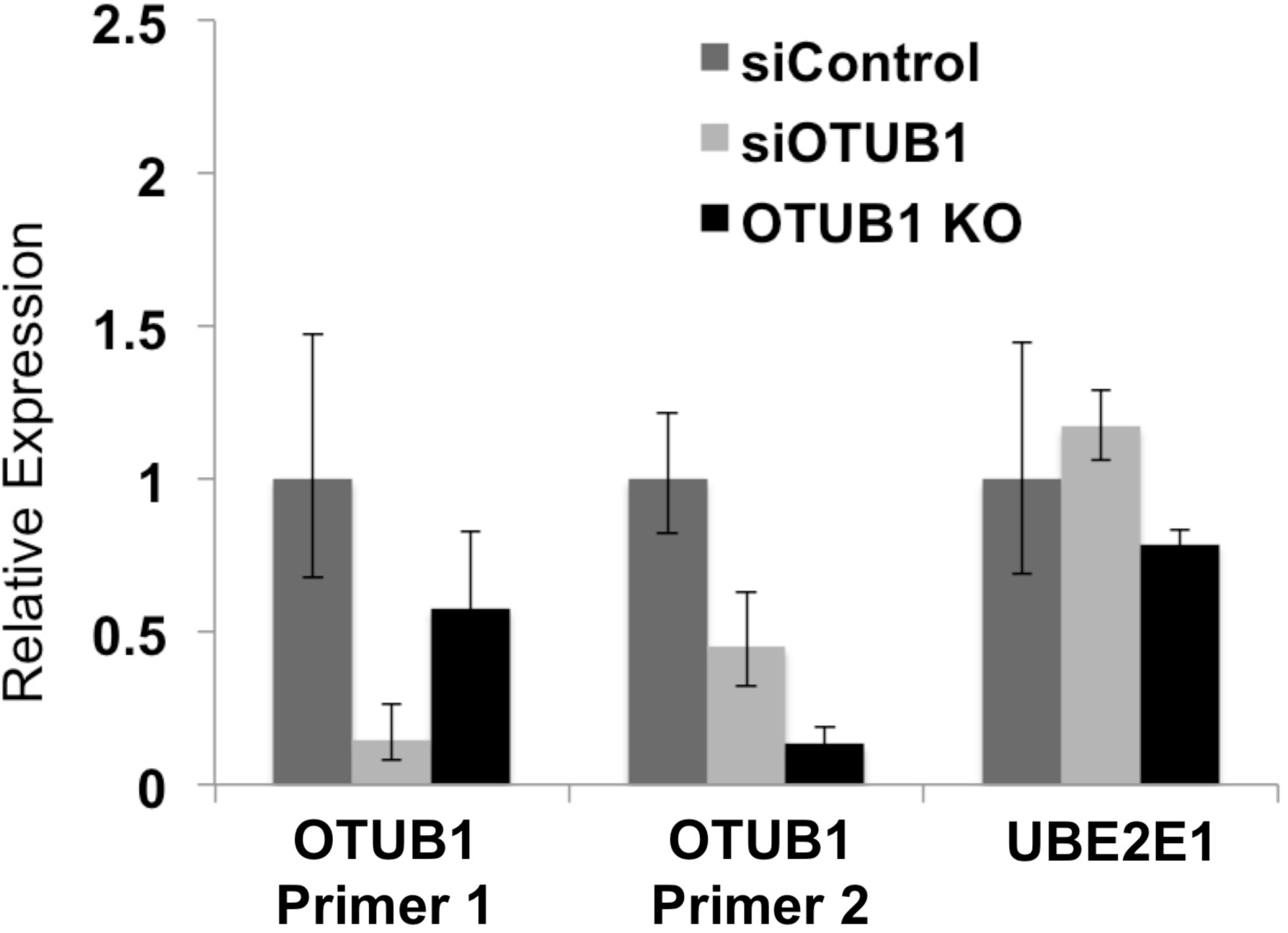
Real-time PCR analysis of OTUB1 and UBE2E1 transcripts in U2OS cells. Total mRNA was extracted from wild-type, siOTUB1, and OTUB1 KO U2OS cells and was quantified by RT-PCR using primers targeting OTUB1 upstream (primer 1), OTUB1 at the CRISPR indel site (primer 2), and UBE2E1. Expression was normalized to TATA-box-binding protein (TBP).

**Figure S4:**
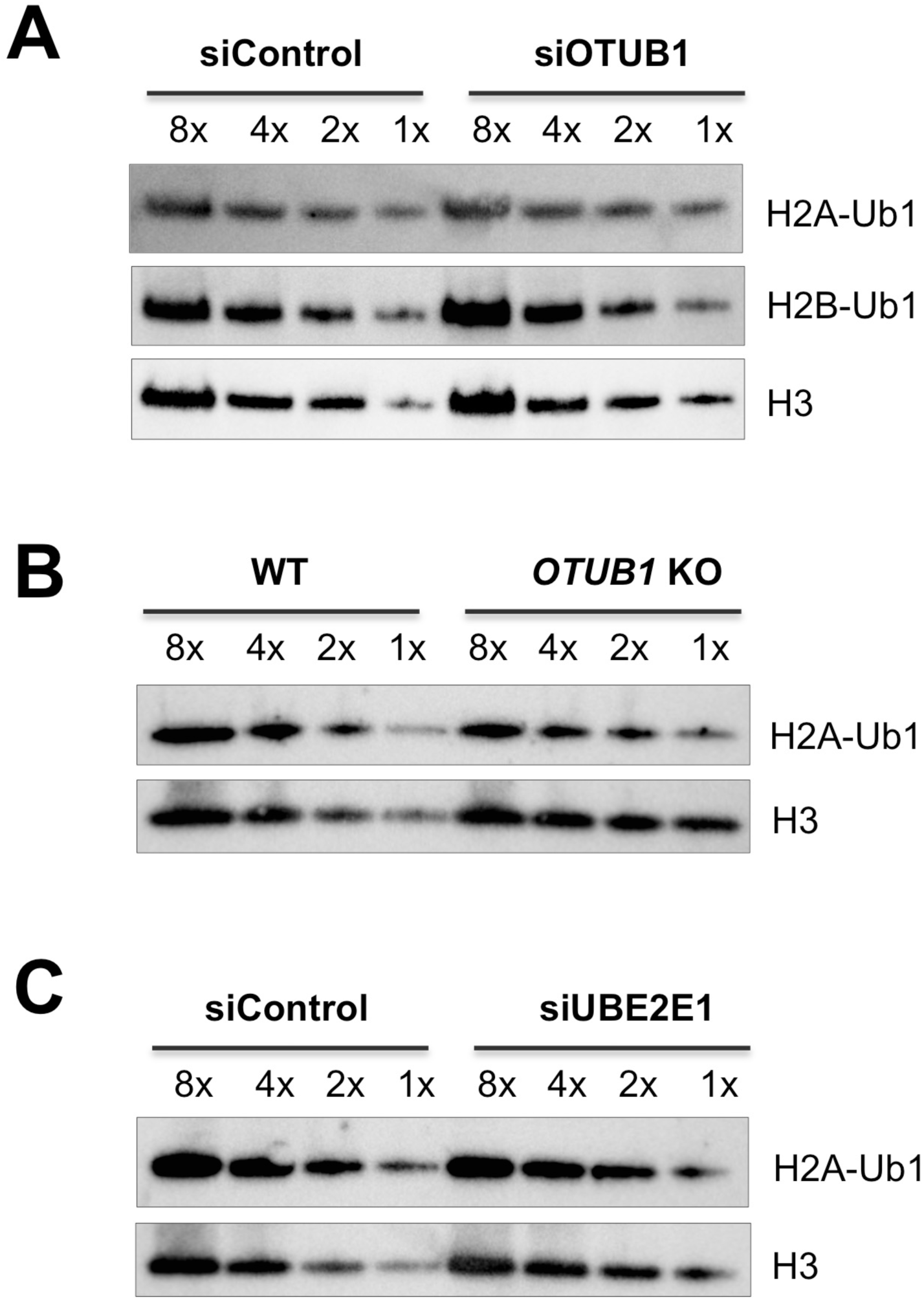
Global levels of monoubiquitinated H2A(K119) in the dynamic range of acid extracted histone from OTUB1 knockdown/knockout and UBE2E1 knockdown U2OS cells. Samples from Figure 6A, 6B and 6C were loaded in gradual two fold dilutions and analyzed by western blotting.

